# Synaptic Ultrastructural Alterations in Human Focal Cortical Dysplasia: Insights from Volume Electron Microscopy

**DOI:** 10.1101/2025.03.05.641779

**Authors:** Gyu Hyun Kim, Na-young Seo, Seung-Ki Kim, Yang Hoon Huh, Ji Yeoun Lee, Kea Joo Lee

## Abstract

Focal cortical dysplasia (FCD) is a developmental disorder of the cerebral cortex and a leading cause of drug-resistant epilepsy in children and young adults. A disrupted excitation-inhibition (E-I) balance is a hallmark of neuronal hyperexcitability in FCD, yet the underlying synaptic ultrastructural alterations remain poorly understood. Using volume electron microscopy, we performed a detailed morphological assessment of synaptic density, size, and organelle distribution within synapses in the temporal cortical layer III of an FCD patient. Our quantitation revealed a reduced density of inhibitory (symmetric) synapses on pyramidal neuron cell bodies. Notably, the dysplastic region displayed a lower density of excitatory (asymmetric) synapses but contained extra-large excitatory synapses, which contained an increased number of synaptic vesicles. Additionally, inhibitory synapses were located further away from the nearest excitatory synapses along distal dendrites, possibly weakening the effectiveness of shunting inhibition in the dysplastic area. There was also a decrease in presynaptic boutons housing mitochondria and in postsynaptic protrusions containing a spine apparatus, pointing to potential deficits in intracellular calcium handling and synaptic plasticity in the epileptogenic area. Moreover, maladaptive myelination was a prominent feature in the dysplastic region. These findings collectively indicate that synaptic architectural modifications may contribute to the neuronal hyperexcitability associated with epilepsy in FCD.

## Introduction

Focal cortical dysplasia (FCD) is a neurodevelopmental disorder of the neocortex characterized by disruptions in neuronal migration and differentiation [1, 2]. It is recognized as the leading cause of drug-resistant epilepsy in both pediatric and adult populations [3, 4]. Due to the poor response of these seizures to anti-epileptic medications, surgical removal of the affected cortical region remains the most effective treatment. The International League Against Epilepsy (ILAE) classifies FCD into three primary subtypes based on histopathological features: Type I, involving abnormal cortical layering without significant cytological abnormalities; Type II, marked by cortical disorganization along with dysmorphic neurons and/or balloon cells; and Type III, which occurs alongside other neuropathological conditions such as hippocampal sclerosis, tumors, or vascular malformations [1, 5-7].

Epilepsy is characterized by excessive synchronized neuronal discharges leading to hyperexcitability [8]. A key mechanism implicated in epileptogenesis is the disruption of the balance between excitatory and inhibitory synaptic activity [9]. While previous electrophysiological and histological studies have provided valuable insights into epilepsy mechanisms in FCD [10-15], the exact cellular alterations responsible for hyperexcitability remain unclear, partly due to limited data on the intricate synaptic architecture of the human cortex.

Immunocytochemical studies using light microscopy have reported a reduced number of basket and chandelier cells—key inhibitory interneurons that target the soma and axon initial segments of pyramidal neurons—in both animal models and human epileptic cortex [8, 15-19]. Consistent with these findings, electrophysiological analysis has demonstrated a reduction in inhibitory postsynaptic currents in pyramidal neurons within dysplastic cortical regions [20], suggesting an overall shift toward increased excitation. However, there exist some conflicting reports regarding glutamate receptor expression: some analyses indicate upregulation of AMPA and NMDA receptor subunits (GluA2/3, GluN1, GluN2A/B) in the dysplastic cortex [15, 21, 22], while others describe a reduction in GluA2/3 immunoreactivity in epileptic neocortex [23]. Similarly, electron microscopic studies have yielded contradictory results, with some reporting an increase in excitatory (asymmetric) synapses in the epileptogenic cortex [24], whereas others have found a decrease in excitatory synapses or dendritic spines—the primary postsynaptic sites of glutamatergic synapses [25, 26]. These inconsistencies suggest that the mechanisms underlying hyperexcitability in FCD involve complex, heterogeneous changes in excitatory synaptic organization.

Given these uncertainties, a precise three-dimensional (3D) ultrastructural analysis of excitatory synapses is essential for elucidating the pathogenesis of epilepsy linked to FCD. In this study, we employed volume electron microscopy to examine local synaptic alterations in human temporal cortex tissue resected from a patient with FCD type I. This subtype, characterized by cortical dyslamination without significant cytological abnormalities, may serve as an ideal model for investigating intrinsic synaptic mechanisms of hyperexcitability, independent of additional confounding pathologies. We focused on cortical layer III, where pyramidal neurons play a key role in corticocortical propagation of neuronal activity [27], and have been previously shown to undergo seizure-associated neuronal remodeling [8, 28, 29]. We hypothesize that synaptic density and morphology in the dysplastic cortex would be altered in a manner favoring excessive excitation. To test this hypothesis, we conducted within-patient comparisons between the epileptic focus and adjacent histologically-normal region. Using serial block-face scanning electron microscopy (SB-SEM) and high-voltage electron microscopy (HVEM), we analyzed synapse types, density, morphology, and intrasynaptic organelle distribution in the dysplastic region, aiming to identify structural changes that may contribute to neuronal hyperexcitability in FCD.

## Materials and Methods

### Tissue Acquisition and Preparation for Microscopy

Surgery was performed after obtaining informed consent, in accordance with the Declaration of Helsinki. All procedures and the use of human tissue were approved in advance by the institutional review board (Ethics Committee, Seoul National University Medical Center, IRB No. H-0507-509-153). Brain tissue samples were collected from a 16-year-old male patient diagnosed with FCD and experiencing intractable epilepsy. The epileptic focus, localized to the left medial temporal lobe as confirmed by MRI (Supplementary Fig. 1), was surgically resected. Seizure activity was confined to the uncus, without involvement of surrounding regions. During the procedure, the dysplastic cortex containing the epileptic focus and a nearby control area was ablated. Histological evaluation was performed on biopsy specimens using hematoxylin and eosin (H&E) staining and immunohistochemistry with an anti-NeuN antibody, enabling neuropathological classification of FCD type. Cortical layer III cell density in the temporal lobe was quantified using ImageJ (NIH).

For volume electron microscopy, tissue samples were immediately immersion-fixed to preserve ultrastructural integrity. Samples were fixed in 2% paraformaldehyde and 2.5% glutaraldehyde in 0.15M cacodylate buffer (pH 7.4) and sectioned into 150-μm slices with a vibratome (Leica VT 1000S). The slices were further dissected into small blocks containing cortical layer III. These blocks were washed in 0.15M cacodylate buffer, post-fixed in 2% osmium tetroxide/1.5% potassium ferrocyanide for 1 hour, incubated in 1% thiocarbohydrazide (Sigma-Aldrich, Cat #223220) for 20 minutes, and treated with 2% osmium tetroxide for 30 minutes. Samples were subsequently incubated in 1% uranyl acetate at 4°C overnight and in lead aspartate at 60°C for 30 minutes. Following serial ethanol dehydration, the specimens were infiltrated with acetone and embedded in 7% (w/v) conductive Epon 812 resin (EMS, Cat #14120) mixed with Ketjen black powder to enhance conductivity, as described previously [30]. Specimens were mounted on metal stubs and polymerized at 60°C for 48 hours. To assess cortical layering, semithin 100-nm sections were stained with toluidine blue and examined under light microscopy.

### SB-SEM Imaging and Analysis

SB-SEM imaging was conducted using a Merlin 3View SEM system (Carl Zeiss Microscopy GmbH, Oberkochen, Germany). Serial images were acquired with a 30-μm aperture, high vacuum, a voltage of 1.5 kV, an image size of 5,000 × 5,000 pixels, a dwell time of 3.5 µs, and an x–y resolution of 9 nm at a nominal section thickness of 50 nm. Stacks of 200 serial images of cortical layer III were obtained from the control and epileptogenic region, covering a volume of 20,250 μm³ per each stack. In the dysplastic region, where cortical layering was less distinct, images were obtained from 550–900 μm beneath the pial surface, corresponding to cortical layer III [25]. Image stacks were processed using ImageJ and Fiji plugins (http://fiji.sc/wiki/index.php/Fiji), with TrakEM2 used for image alignment.

Spiny dendritic segments, averaging 15.94 ± 3.56 μm in length, were randomly selected. A total of 20 dendritic branches (10 per condition) were manually reconstructed using open-source reconstruction software (https://synapseweb.clm.utexas.edu/) by annotators blinded to experimental conditions. Quantification of spine density, presence or absence of presynaptic boutons, and surface area of postsynaptic density (PSD) was performed. Synapses were classified as asymmetric or symmetric based on PSD thickness [31]. Additional analyses included spine volume, length, spine head and neck diameters, and classification of extra-large synapses (top 20th percentile in PSD area and spine volume), based on prior studies of activity-dependent synaptic structural enlargement [32, 33]. To measure the distance between inhibitory and excitatory synapses, their locations were mapped onto the skeletonized dendrite. Dendritic shaft thickness was measured using maximal and minimal diameters, with a regularity index calculated by dividing the minimal by the maximal diameter. Presynaptic boutons were categorized by mitochondrial presence, and the number of mitochondria was counted per bouton. The presence of a spine apparatus (SA) was determined when at least three smooth endoplasmic reticulum stacks were visible.

### ATUM-SEM imaging and Analysis

For 3D analysis of symmetric synapses on pyramidal neuron somata, we used ATUM-SEM (<u>A</u>utomated <u>T</u>ape-Collecting <u>U</u>ltra<u>m</u>icrotome combined with SEM). Tissue blocks were sectioned into 50 nm ultrathin sections using an ultra-Maxi knife (DiATOME, Biel, Switzerland). Serial sections were collected onto plasma-hydrophilized carbon nanotube-coated PET tapes (Boeckeler Instruments) with an ATUMtome (Boeckeler Instruments Inc., Tucson, USA). The tapes were mounted on a silicon wafer, secured with conductive adhesive tape (Ted Pella), and carbon-coated to prevent charging during SEM imaging. Imaging was performed using a Gemini 300 SEM (Carl Zeiss Microscopy GmbH, Oberkochen, Germany) equipped with an In-lens secondary electron detector or a backscattered electron detector (BSD). Large-area imaging was conducted using Atlas 5 software (Fibics Incorporated, Ottawa, Canada) at 5 kV beam voltage with BSD detection and a 7 µs dwell time. High-resolution images (5 nm per pixel) were obtained as 2 × 2 tiled images, stitched together to generate a composite 90 × 90 μm² image. Image stacks were aligned using the FIJI TrakEM2 plugin. Twelve cortical neurons per group were manually segmented, and the number of presynaptic boutons forming symmetric synapses on somata was quantified. Myelin sheath thickness was assessed by measuring the G-ratio, calculated as the ratio of axonal radius (r) to total nerve fiber radius (R) including both axon and myelin sheath.

### Electron Tomography

To visualize synaptic ultrastructure in cortical layer III, 250 nm-thick sections were collected on 200-mesh grids (G200N, Gilder Grids) for electron tomography using a Bio-HVEM (JEM-1000BEF, JEOL, Tokyo, Japan) operating at 1,000 kV (Korea Basic Science Institute, Ochang, Korea). Excitatory synapses were randomly selected for imaging, and samples were tilted from +60° to −60° in 2° increments, producing a total of 61 tilt images per synapse captured with TEM Recorder software (JEOL System Technology, Tokyo, Japan). The tilt series was aligned and reconstructed into 3D tomograms using Composer and Visualizer-Kai software (TEMography.com, Frontiers Inc., Tokyo, Japan). Virtual slices were extracted, and synaptic vesicles (SVs) were traced using reconstruction software (https://synapseweb.clm.utexas.edu/software-0). To prevent overcounting, only vesicles with clearly defined circular profiles were quantified in serial tilt images. Vesicle distribution, pre- and postsynaptic site volumes, and docked vesicle counts (within 50 nm of the active zone) were analyzed as correlates of synaptic strength.

### Statistics

All data are presented as mean ± SEM. Statistical analyses were performed using GraphPad Prism software (RRID: SCR_002798). Synaptic density measurements from SB-SEM data were treated as single observations per dendrite. Normality was assessed, and comparisons between groups were conducted using either an unpaired Student’s t-test (for normally distributed data) or a Mann–Whitney U test (for non-normally distributed data). Statistical significance was set at P < 0.05.

## Results

### Histological Analysis

Tissue samples were obtained from a patient who underwent epilepsy surgery (Supplementary Fig. 1a-e). Compared to the control cortex, disorganized cortical lamination was observed in the epileptogenic region using H&E staining. Although cell density within cortical layer III (550–900 μm from the pial surface) was reduced in the epileptogenic area, dysmorphic neurons or balloon cells were not identified (Supplementary Fig. 1f-h). Immunohistochemistry with an anti-NeuN antibody also highlighted a characteristic radial microcolumnar arrangement (Supplementary Fig. 1i), a predominant feature of FCD [34]. Based on these histopathological observations, the patient was diagnosed with FCD type I.

### Ultrastructural Synaptic Analysis of in Cortical Layer III pyramidal neurons

Given the well-established E-I imbalance in FCD, we investigated whether synaptic alterations favoring hyperexcitation might involve changes in excitatory and/or inhibitory synapses on cortical pyramidal neurons. We first assessed inhibitory synapses on pyramidal neuron somata, primarily targeted by parvalbumin-expressing interneurons that exert strong somatic inhibition [35]. Quantitative analysis of ATUM-SEM datasets revealed a significant reduction in the density of inhibitory synapses on pyramidal cell bodies in the epileptogenic region compared to control tissue (Supplementary Fig. 2) (synapses/100 μm^3^: control, 1.86 ± 0.32; epileptogenic, 0.94 ± 0.24; p = 0.029).

To further examine excitatory synaptic changes, we analyzed dendritic spines, which serve as the primary sites of excitatory synapses in pyramidal neurons. Using SB-SEM, 3D reconstructions of dendritic segments were generated from both control and epileptogenic regions (Fig. 1a; Supplementary Fig. 3). Analysis of twenty reconstructed dendritic segments (mean segment length: 15.94 ± 3.56 μm) demonstrated a significant reduction in spine density in the epileptogenic region, affecting both spines with and without presynaptic partners (Fig. 1b) (total spines/10 μm: control, 11.8 ± 1.09; epileptogenic, 4.3 ± 0.31; p < 0.0001; synaptic spines/10 μm: control, 8.4 ± 0.95; epileptogenic, 2.9 ± 0.26; p < 0.0001; non-synaptic spines/10 μm: control, 3.3 ± 0.65; epileptogenic, 1.3 ± 0.40; p = 0.016). Interestingly, despite this overall spine loss, some remaining spines in the epileptogenic region exhibited notably larger volumes, while spine length remained unchanged (Supplementary Fig. 4a–d) (spine volume [μm^3^]: control, 0.15 ± 0.01; epileptogenic, 0.35 ± 0.05; p = 0.048; spine length [μm]: control, 1.82 ± 0.06; epileptogenic, 1.70 ± 0.10; p = 0.26). Moreover, the PSD areas of the largest 20% of spines were significantly enlarged in the epileptogenic region (Supplementary Fig. 4e–f) (PSD area [μm^2^]: control, 0.91 ± 0.06; epileptogenic, 1.40 ± 0.24; p = 0.013).

**Fig. 1.**
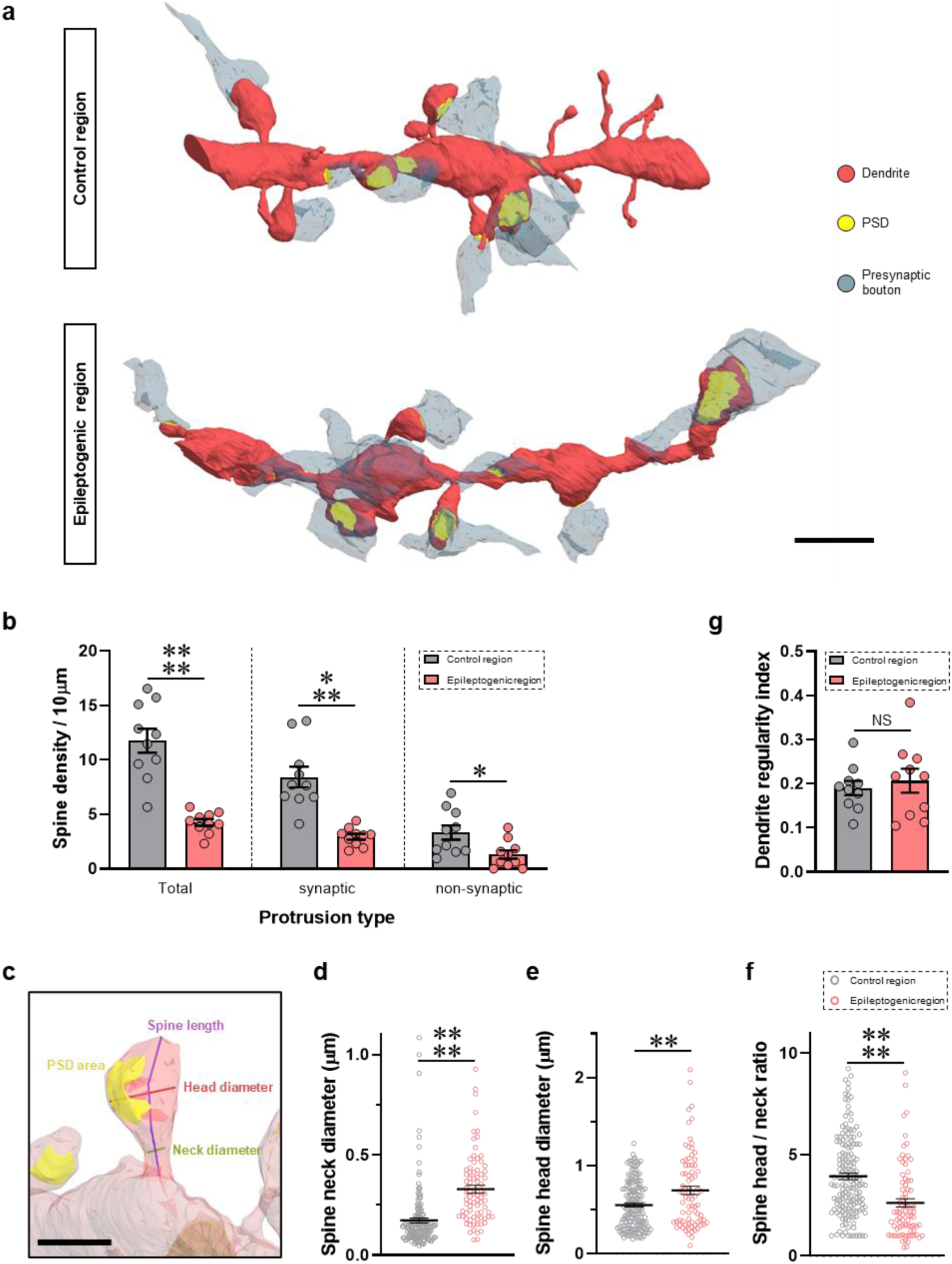
Reduced spine density and abnormal spine enlargement in the epileptic cortex. **a** 3D reconstruction of SB-SEM images showing dendritic segments with PSDs and presynaptic boutons in cortical layer III of the control and epileptogenic cortex. Red, dendrite; yellow, PSD; light blue, presynaptic bouton. Scale bar, 2 µm. **b** Quantification of spine density along the dendrite in the control and epileptogenic regions (n = 10 dendrites). ****P<0.0001; ***P<0.001; *P<0.05; unpaired Student’s t-test. **c** Morphological analysis of individual spine properties. Yellow, PSD; blue line, spine length; red line, spine head diameter; green line, spine neck diameter. Scale bar, 1 µm. **d-f** Quantification of spine neck diameters (d), spine head diameters (e), and the ratio of spine head diameter to neck diameter (f) (control, n = 156; epilepsy, n = 79). ****P<0.0001; **P<0.01; Mann-Whitney U test. **g** Quantification of dendritic regularity in the control and epileptogenic regions (n = 10 dendrites). NS, not significant. unpaired Student’s t-test. Data are represented as mean ± SEM.

Further morphological analysis revealed increased diameters of both spine heads and necks in the epileptogenic region (Fig. 1c–e) (spine head [μm]: control, 0.55 ± 0.02; epileptogenic, 0.72 ± 0.05; p = 0.007; spine neck [μm]: control, 0.15 ± 0.01; epileptogenic, 0.30 ± 0.02; p < 0.0001). The spine head-to-neck diameter ratio was significantly reduced in the dysplastic area (Fig. 1f) (control, 3.92 ± 0.16; epileptogenic, 2.61 ± 0.21; p < 0.0001), suggesting that spine necks expanded disproportionately relative to their heads. This structural alteration may reduce the electrochemical compartmentalization of synaptic signals, facilitating the spread of excitatory activity and contributing to hyperexcitability. Additionally, we observed varicose swelling in both epileptogenic and control dendrites, a feature commonly linked to excitotoxic damage or seizure-related remodeling [36, 37]. However, quantitative assessment of dendritic regularity showed no significant differences between groups (Fig. 1g) (control, 0.19 ± 0.02; epileptogenic, 0.21 ± 0.03; p = 0.60).

To confirm the observed reduction in spine density and assess inhibitory synapses on distal dendrites, serial EM images were analyzed, quantifying excitatory (asymmetric) and inhibitory (symmetric) synapses on dendritic shafts and spines (Fig. 2a). Again, asymmetric synapse density was significantly lower in the epileptogenic region, whereas symmetric synapse density remained unchanged (Fig. 2b–c) (asymmetric synapses/10 μm: control, 9.7 ± 1.04; epileptogenic, 4.4 ± 0.30; p = 0.0001; symmetric synapses/10 μm: control, 0.7 ± 0.26; epileptogenic, 0.7 ± 0.20; p = 0.86).

**Fig. 2.**
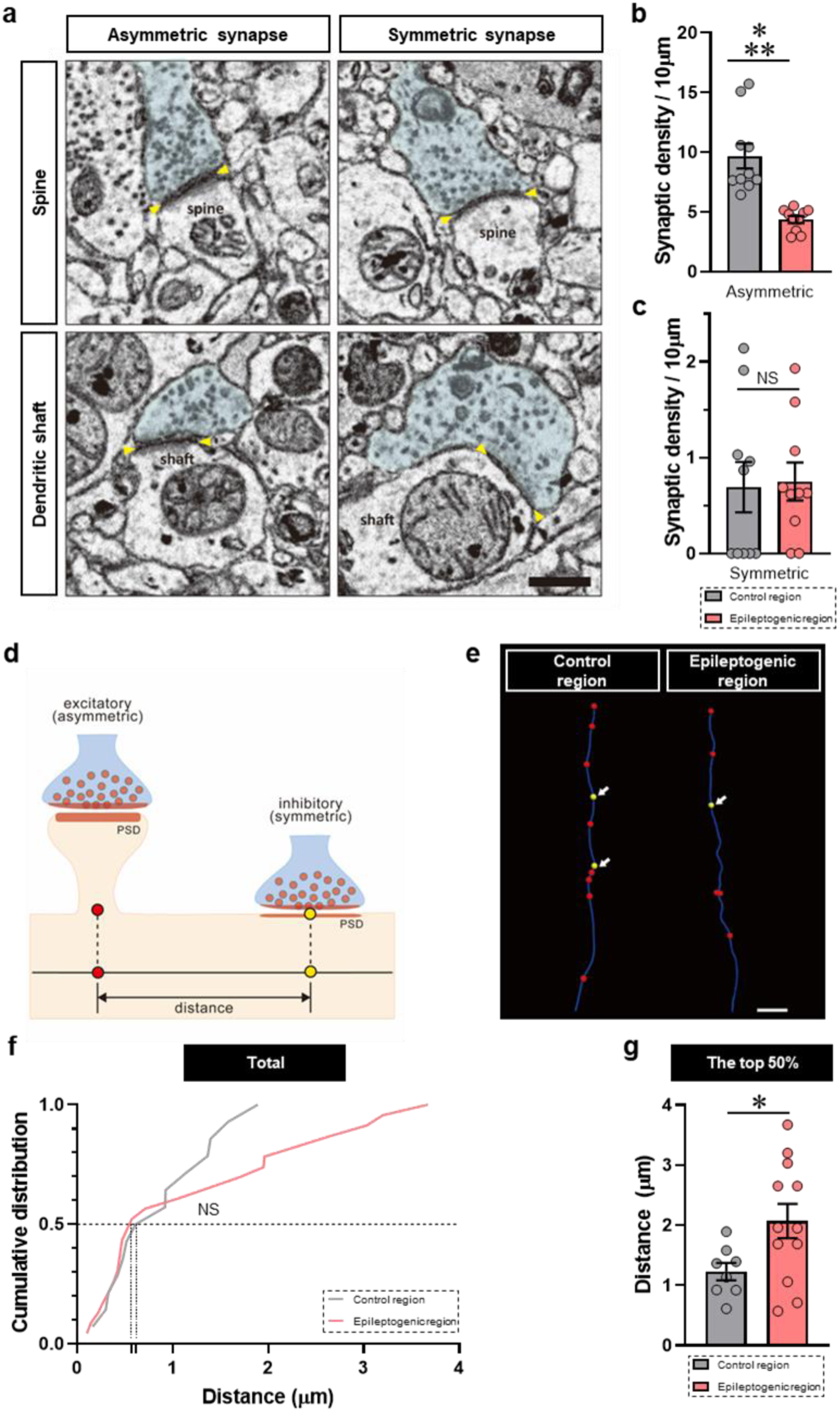
Spatial distribution of excitatory and inhibitory synapses along distal dendrites. **a** Representative EM images of asymmetric (excitatory) and symmetric (inhibitory) synapses formed on dendritic spines or directly onto the shaft. Yellow arrow, PSD. Scale bar, 0.5 µm. **b**, **c** Quantification of asymmetric (b) and symmetric (c) synaptic density (n = 10 dendrites). NS, not significant. ***P<0.001; unpaired Student’s t-test. **d** Schematic illustration of measuring the distance between excitatory and inhibitory synapses. Yellow circle, inhibitory synapse; red circle, excitatory synapse. **e** Representative image of skeletonized dendritic traces with annotated synapses. The distance from an inhibitory synapse (white arrow) to the nearest excitatory synapse was measured. Scale bar, 1 µm. **f** Cumulative distribution of distance measurements between inhibitory and excitatory synapses (control, n = 14; epilepsy, n = 23). NS, not significant; Kolmogorov-Smirnov test. **g** Average distance between inhibitory and excitatory synapses in the top 50% of the population in F (control, n = 8; epilepsy, n = 12). *P<0.05; unpaired Student’s t-test. Data are represented as mean ± SEM.

### Spatial Distribution of Excitatory and Inhibitory Synapses

The precise spatial arrangement of excitatory and inhibitory synapses is crucial for shunting inhibition, which limits excessive excitatory input [38, 39]. We hypothesized that increased separation between inhibitory and excitatory synapses in distal dendrites might impair local inhibition, thereby contributing to seizure activity. To test this, inhibitory synapse locations were mapped on skeletonized dendrite traces, and the distance to the nearest excitatory synapse was measured (Fig. 2d–e). In the epileptogenic region, cumulative distribution analysis revealed a greater proportion of excitatory synapses positioned ≥0.6 μm from inhibitory synapses (Fig. 2f–g) (cumulative distribution of distance between inhibitory and excitatory synapses [μm]: control, 0.86 ± 0.15; epileptogenic, 1.24 ± 0.24; p = 0.34; distance of top 50% [μm]: control, 1.23 ± 0.15; epileptogenic, 2.07 ± 0.29; p = 0.027). This suggests that inhibitory inputs in the epileptogenic region may be less effective in suppressing local excitatory activity, increasing the potential for hyperexcitability.

### Intrasynaptic Organelle Analysis (Vesicles, Mitochondria, and Spine Apparatus)

To explore whether SV organization was altered in enlarged excitatory synapses within the epileptogenic region, we performed electron tomography using a HVEM. Each synapse was imaged across 61 tilt angles (±60° in 2° increments) (Fig. 3a; Supplementary Movie 1). A total of 26 presynaptic boutons (12 control, 14 epileptogenic) were reconstructed, allowing segmentation of synaptic components including postsynaptic spines, presynaptic boutons, presynaptic mitochondria, and SVs (Fig. 3b). In agreement with SB-SEM findings, spine head volume was significantly increased in the epileptogenic region (Fig. 3c), as was presynaptic bouton volume (Fig. 3d) (spine head volume [μm^3^]: control, 0.06 ± 0.01; epileptogenic, 0.12 ± 0.03; p = 0.031; presynaptic bouton volume [μm^3^]: control, 0.15 ± 0.02; epileptogenic, 0.24 ± 0.03; p = 0.034).

**Fig. 3.**
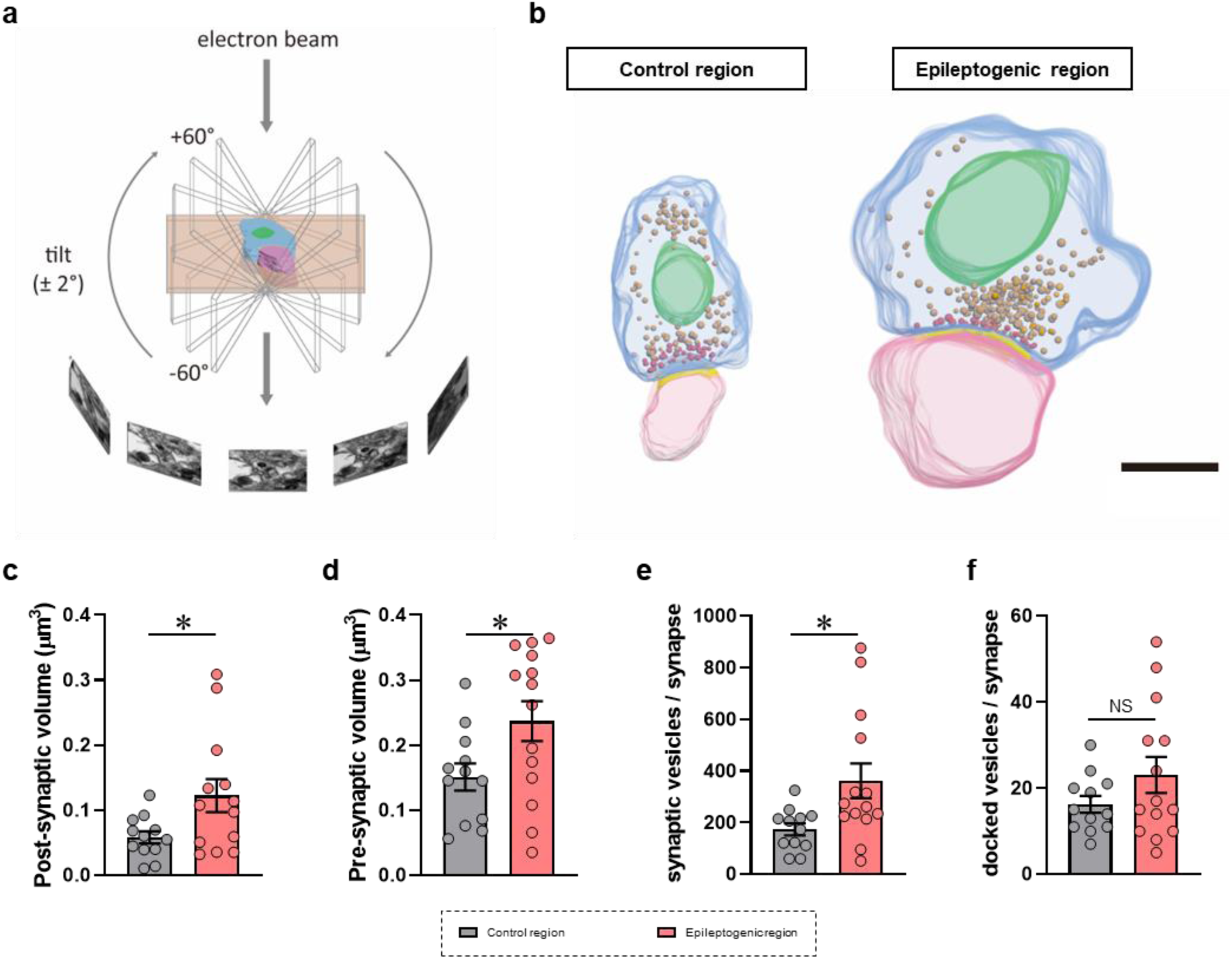
Enlarged excitatory synapses with increased vesicles in the epileptogenic area. **a** Schematic representation of a synapse imaged using high-voltage EM tilt-series. Each series consists of 61 images captured every 2° from -60° to +60°. **b** Representative 3D reconstructions of excitatory synapses, showing the spine head (pink), PSD (yellow), and presynaptic bouton (light blue) containing docked vesicles (red), reserve pool vesicles (orange), and mitochondria (green). Scale bar, 0.5 µm. **c**, **d** Quantification of postsynaptic (c) and presynaptic (d) volume (n = 12-14 synapses per region). *P<0.05; unpaired Student’s t-test. **e**, **f** Quantification of total (e) and docked (f) synaptic vesicles in presynaptic boutons (n = 12-14 boutons per region). NS, not significant. *P<0.05; unpaired Student’s t-test. Data are represented as mean ± SEM.

To assess SV distribution, we quantified the total SV pool and the number of docked vesicles (Fig. 3b, e–f). Docked vesicles—those in direct contact with the active zone—were similar between groups. However, the total number of SVs was significantly higher in epileptogenic boutons than in control boutons (Fig. 3e–f; Supplementary Fig. 5) (total SVs/bouton: control, 173.8 ± 23.2; epileptogenic, 361.5 ± 67.5; p = 0.042; docked SVs/bouton: control, 16.3 ± 1.9; epileptogenic, 23.1 ± 4.2; p = 0.38), suggesting an increased vesicle reserve that could enhance neurotransmitter release and contribute to a prolonged state of neuronal hyperactivity.

Presynaptic mitochondria are essential for supporting synaptic transmission, regulating calcium dynamics, and sustaining energy homeostasis [40, 41]. We next examined whether presynaptic mitochondrial distribution was altered in the epileptogenic region (Fig. 4). Although the proportion of boutons lacking mitochondria did not differ significantly between groups, the density of mitochondria-containing boutons was significantly lower in the epileptogenic region (Fig. 4b) (boutons without mitochondria/10 μm: control, 3.8 ± 1.04; epileptogenic, 1.9 ± 0.33; p = 0.11; boutons with mitochondria/10 μm: control, 6.6 ± 0.41; epileptogenic, 3.3 ± 0.31; p < 0.0001). Notably, both mitochondrial number and volume per bouton were greater in the epileptogenic region compared to the control (Fig. 4c–d) (mitochondria number/bouton: control, 1.2 ± 0.06; epileptogenic, 1.7 ± 0.15; p = 0.03; individual mitochondria volume [μm^3^]: control, 0.08 ± 0.01; epileptogenic, 0.15 ± 0.01; p < 0.0001), potentially reflecting compensatory mitochondrial redistribution in response to synaptic remodeling.

**Fig. 4.**
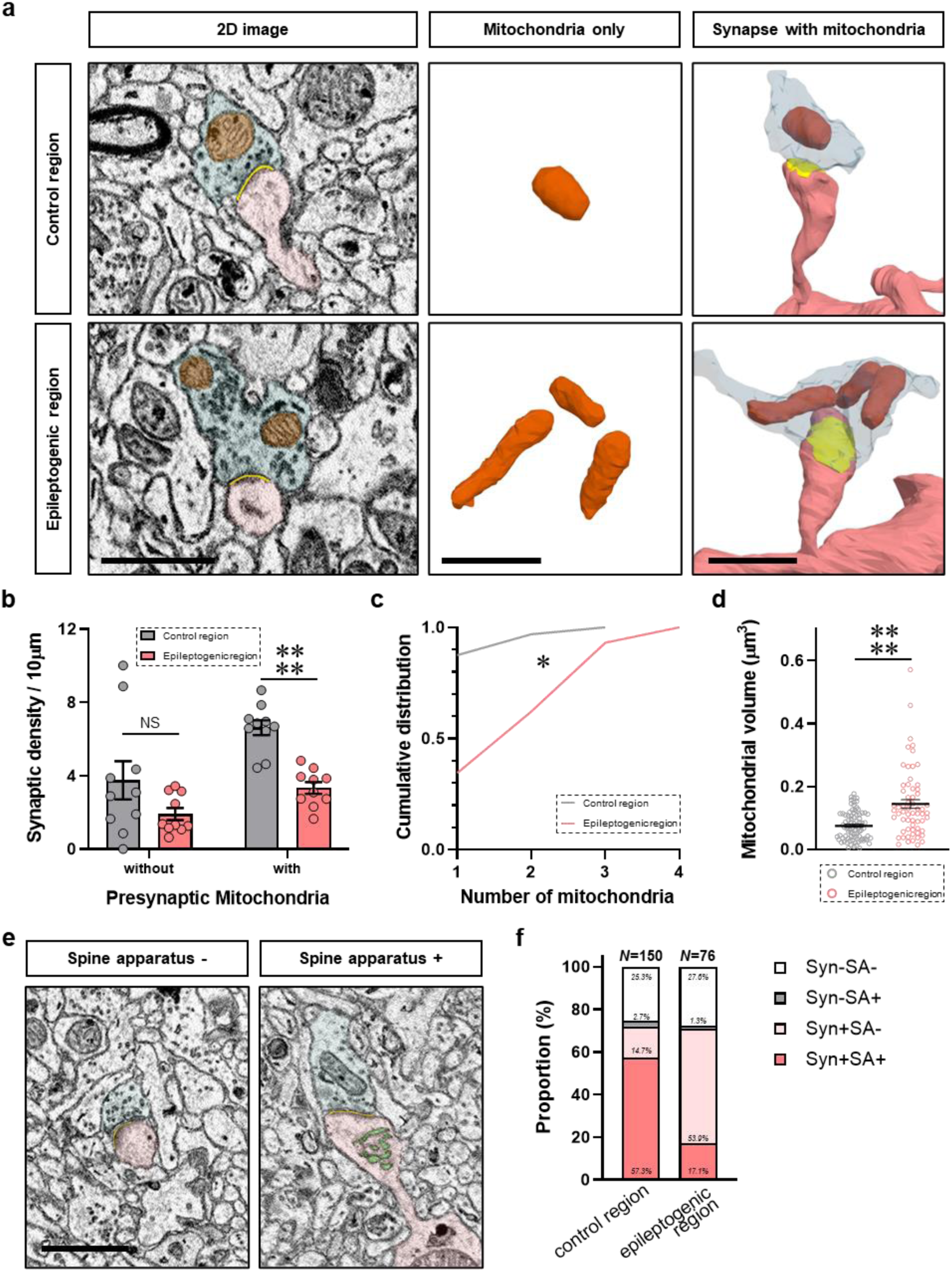
Altered presynaptic mitochondria and spine apparatus in the epileptogenic cortex. **a** Representative EM images of synapses containing presynaptic mitochondria (left panels). 3D reconstructions of presynaptic mitochondria and synaptic components (middle and right panels, respectively). Light blue, presynaptic bouton; orange, mitochondria; yellow, PSD; red, spine. Scale bar, 1 µm. **b** Quantification of synapse density based on the presence or absence of presynaptic mitochondria in the control and epileptogenic cortical regions (n = 10 dendrites). NS, not significant. ****P<0.0001; unpaired Student’s t-test. **c** Cumulative distribution of mitochondrial numbers per bouton (control, n = 64; epilepsy, n = 35). *P<0.05; Kolmogorov-Smirnov test. **d** Average volume of individual presynaptic mitochondria (control, n = 82; epilepsy, n = 62). ****P < 0.0001; Mann-Whitney U test. **e** Representative EM images of postsynaptic protrusions with and without spine apparatus. Scale bar, 1 µm. **f** Percentage of synaptic and non-synaptic protrusions containing spine apparatus (n = 8-10 dendrites/region). NS, not significant; Mann-Whitney U test for non-synaptic protrusion. ****P<0.0001; unpaired Student’s t-test for synaptic and total protrusions. Data are represented as mean ± SEM.

The SA, a specialized organelle derived from the smooth endoplasmic reticulum, regulates synaptic calcium dynamics and plasticity [42, 43]. Given the presence of abnormally large spines in the epileptogenic region, we examined whether these structures contained the SA (Fig. 4e–f). While most spines in the control region exhibited an SA, a substantial proportion of enlarged spines in the epileptogenic region lacked this organelle (Fig. 4f) (synaptic protrusions with SA [%]: control, 84.6 ± 6.0; epileptogenic, 24.0 ± 8.4; p < 0.0001; non-synaptic protrusions with SA [%]: control, 12.0 ± 7.1; epileptogenic, 6.3 ± 5.6; p = 0.51). This suggests that these hypertrophic spines may be deficient in key organelles required for calcium signaling and synaptic plasticity.

### Myelination Changes in the Epileptogenic Cortex

Neuronal activity modulates myelination, which is essential for network synchronization and efficient signal conduction [44, 45]. A recent study indicates that aberrant neuronal activity can induce maladaptive myelination, potentially exacerbating seizure susceptibility and progression [46]. To evaluate axonal myelination changes in FCD, we measured the G-ratio (the ratio of inner axonal diameter to total fiber diameter, including the myelin sheath) in control and epileptogenic regions (Fig. 5). Our analysis revealed a significant reduction in G-ratio values in the epileptogenic region, indicating increased myelin thickness (G-ratio: control, 0.731 ± 0.012; epileptogenic, 0.691 ± 0.013; p = 0.028). This suggests that hyperactivity-induced myelination changes may contribute to altered neuronal network properties in FCD.

**Fig. 5.**
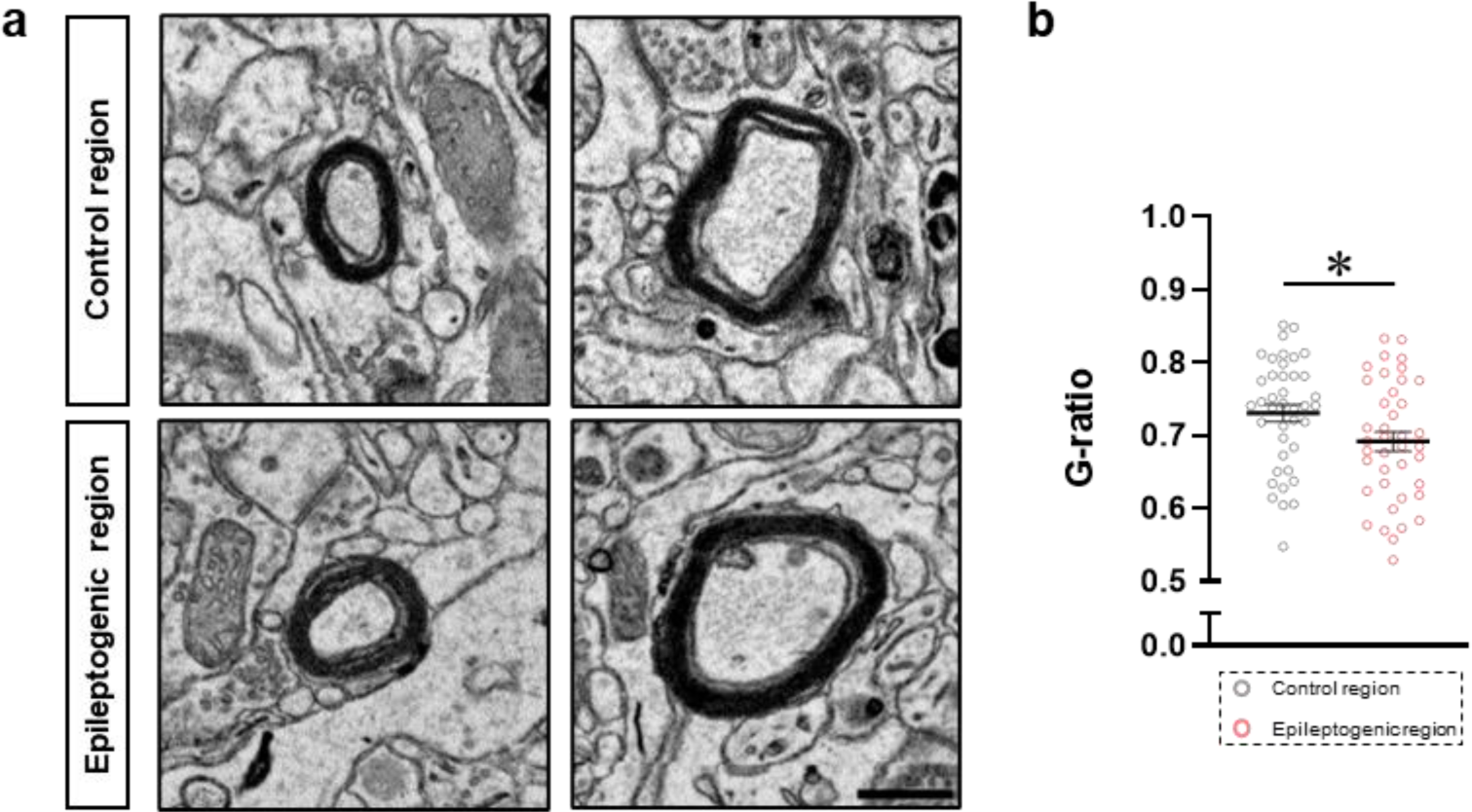
Maladaptive myelination in the epileptogenic cortex. **a** Representative EM images of myelinated axons in the control and epileptogenic areas. Scale bar, 0.5 µm. **b** Quantification of G-ratio values (inner axonal diameter divided by total fiber diameter, including the axon and myelin sheath) in control and epileptogenic regions (n = 40 per group). *P<0.05; unpaired Student’s t-test. Data are represented as mean ± SEM.

## Discussion

FCD characterized by disrupted cortical structure is a leading cause of drug-resistant epilepsy, particularly in pediatric and young adult populations [3, 4]. Although genetic mutations affecting the mTOR signaling pathway, environmental influences, and somatic mosaic mutations have been associated with FCD [47-49], the exact mechanisms underlying abnormal cortical network formation remain largely unclear. To investigate the neuroanatomical correlates of neuronal hyperexcitation, we utilized advanced volume electron microscopy techniques, including SB-SEM, ATUM-SEM, and electron tomography, to analyze ultrastructural synaptic alterations in the epileptogenic cortex of a patient with FCD Type I. Our findings provide novel insights into local synaptic and axonal alterations that may contribute to cortical hyperexcitability and seizure propagation in FCD. The observed changes in inhibitory and excitatory synapses on layer III pyramidal neurons suggest that multiple mechanisms underlie seizure susceptibility in FCD (Supplementary Fig. 6).

Histological examination confirmed characteristic features of FCD type I, such as disrupted cortical layering and a distinct radial microcolumnar arrangement, with no evidence of dysmorphic neurons or balloon cells (Supplementary Fig. 1) [34]. Additionally, a reduction in cell density within cortical layer III was observed, which may reflect aberrant neuronal distribution or seizure-related neuronal loss, ultimately influencing the E-I balance.

At the synaptic level, we identified a decline in inhibitory synapses on the soma of layer III pyramidal neurons within the epileptogenic cortex (Supplementary Fig. 2). Since somatic inhibition, predominantly mediated by parvalbumin-positive interneurons, plays a crucial role in regulating pyramidal neuron excitability [35], this reduction likely contributes to increased cortical excitability. Our findings are consistent with previous studies reporting a selective loss of parvalbumin-expressing interneurons and diminished inhibitory postsynaptic currents in pyramidal neurons within FCD-affected cortex [8, 15, 20], reinforcing the notion of an excitation-dominant state. Somewhat paradoxically, despite the hyperexcitable nature of the epileptogenic cortex, we observed a marked reduction in dendritic spine density in the distal dendrites of pyramidal neurons (Fig. 1). This aligns with prior research demonstrating a decrease in excitatory synapses or dendritic spine loss in cortical regions of epileptic patients [25, 37, 50]. Further morphological analysis of the remaining dendritic spines in the epileptogenic cortex revealed that a subset displayed significantly enlarged spine heads and necks (Fig. 1; Supplementary Fig. 4), along with an expanded PSD area. Notably, the constriction of spine necks was less pronounced in the epileptogenic cortex, potentially impairing electrochemical compartmentalization between spine heads and dendritic shafts. Such alterations may facilitate greater signal diffusion within the dendritic arbor, thus exacerbating neuronal excitability. This observation reminds me of recent findings in mouse models of schizophrenia, where an overabundance of extra-large synapses resulted in supralinear dendritic and somatic integration, leading to an increase in neuronal firing [51]. Given that layer III pyramidal neurons serve as primary mediators of cortico-cortical communication, pathological amplification of excitatory synaptic activity in these neurons may enhance aberrant network dynamics and facilitate widespread seizure propagation through hyperactive cortico-cortical circuits.

Analysis of excitatory and inhibitory synapse distribution on distal dendrites revealed a selective loss of asymmetric (excitatory) synapses, while symmetric (inhibitory) synapses—primarily innervated by somatostatin-expressing interneurons—remained unaffected (Fig. 2). Consequently, inhibitory and excitatory synapses were positioned farther apart on the same dendritic shafts in the epileptogenic region. Given the critical role of synaptic spatial arrangements in shunt inhibition [38, 39], this increased synapse spacing suggests that inhibitory synapses may be less effective at counteracting excitatory input, further contributing to cortical hyperexcitability. Since precise inhibitory-excitatory synapse positioning is essential for dendritic integration, neural computation, and overall network stability [52], we propose that alterations in this spatial arrangement in pyramidal neuron dendrites contribute to both hyperexcitability and impaired synaptic plasticity in FCD patients.

Electron tomography confirmed that excitatory synapses in the epileptogenic region contained an increased number of SVs within presynaptic terminals, despite no change in the number of docked vesicles (Fig. 3; Supplementary Fig. 5). This accumulation of SVs suggests potential disruptions in vesicle cycling or prolonged neurotransmitter release machinery, which could enhance synaptic efficacy and promote excessive excitation. Additionally, the reduced presynaptic boutons in the epileptogenic cortex exhibited a greater number of mitochondria, which were also significantly enlarged (Fig. 4). This suggests possible compensatory mitochondrial redistribution or remodeling to maintain calcium homeostasis and ATP production for heightened synaptic transmission—an adaptive response observed in other hyperexcitable neuronal conditions [40, 41]. In parallel to presynaptic mitochondria, the SA in a subpopulation of spines is involved in calcium homeostasis and synaptic plasticity [42, 43]. The SA was notably absent in a significant proportion of enlarged spines within the epileptogenic cortex (Fig. 4). This may further exacerbate synaptic dysfunction, potentially contributing to seizure-related synaptic remodeling and impaired synaptic plasticity.

Additionally, examination of axonal myelination revealed a significant reduction in G-ratio values (Fig. 5), indicative of increased myelin sheath thickness in the epileptogenic cortex. Since activity-dependent myelination is crucial for synchronizing neuronal networks and modulating seizure activity [44, 45], aberrant neuronal activity may drive maladaptive myelination, further influencing seizure propagation through altered conduction velocities and network connectivity [46]. These findings suggest that hyperactivity-induced changes in myelination could be an additional factor contributing to the pathophysiology of FCD.

Although our 3D EM data were obtained from a significantly larger cortical volume than in previous conventional transmission EM work, they are derived from a single case, limiting their extrapolation to the broader population of FCD patients with epilepsy. Further studies are required to clarify how synaptic connectivity is altered in specific neuron types and circuit levels, including interactions with glia cells, and whether these changes are consistently observed across different FCD cases.

From a methodological standpoint, tissue fixation plays a critical role in preserving ultrastructure for high-resolution electron microscopy, ensuring accurate morphological analysis at the synaptic and subcellular levels. While perfusion fixation is widely regarded as the gold standard for brain tissue preservation due to its ability to rapidly and uniformly fix tissues in vivo, its application is not always feasible in clinical settings, particularly for human surgical specimens. In this study, immersion fixation was employed, and despite its potential limitations, the cortical tissue exhibited excellent ultrastructural preservation (Fig. 2, 4–5; Supplementary Movie 1). This allowed for a detailed and quantitative assessment of synaptic morphology and intracellular organelles, underscoring the feasibility of utilizing surgically resected and immersion-fixed human brain tissue for high-resolution ultrastructural investigations, thereby enabling further research into epilepsy-associated microstructural alterations and potentially broadening the scope of neuropathological studies on resected tissue samples.

While anti-epileptic drugs could potentially influence synaptic architecture, their effects can be ruled out in this study, as both control and dysplastic cortical regions were obtained from the same patient undergoing identical treatment. Thus, the observed differences in synaptic organization are more likely due to FCD-related pathological changes. However, it remains possible that some alterations in the epileptogenic cortex result from compensatory mechanisms or secondary responses aimed at mitigating epilepsy. Future studies integrating electrophysiological recordings and computational simulation modeling would be invaluable for determining the functional impact of these synaptic ultrastructural changes in the FCD cortex.

## Conclusion

In conclusion, our study using volume electron microscopy provides novel insights into the synaptic and axonal abnormalities in FCD type I, highlighting ultrastructural alterations that may contribute to cortical hyperexcitability. Key findings include a reduced number of inhibitory synapses on pyramidal neuron somata, the presence of extra-large excitatory synapses with increased presynaptic vesicles, an elongated distance between inhibitory and excitatory synapses, and maladaptive myelination. Together, these structural alterations suggest that multiple mechanisms underlie hyperexcitability and seizure propagation in FCD. Further studies are needed to determine the functional impact of these changes and to explore potential therapeutic strategies for restoring synaptic balance in epilepsy linked to FCD.

## Abbreviations

ATUM: Automated tape-collecting ultramicrotome
BSD: backscattered electron detector
E-I: Excitation-Inhibition
FCD: Focal cortical dysplasia
H&E: hematoxylin and eosin
HVEM: High-voltage electron microscopy
ILAE: International League Against Epilepsy
PSD: Postsynaptic density
SA: Spine apparatus
SV: Synaptic vesicles
SB-SEM: Serial block-face scanning electron microscopy
3D: three-dimensional

## Supplementary information

The online version contains supplementary material available at https://doi.org~~~~

## Acknowledgments

The authors thank Dr. Namhee Kim (Rush University) for statistical advice and Dr. Sang Hoon Lee (KBRI imaging facility) for technical assistance with electron microscopy and Mr. Chan Hee Lee and Ms. Ji Won Shin, Na Young Do, and Meeji Kim for image segmentation.

## Author contributions

K.J.L., J.Y.L, and Y.H.H. designed the experiments, and supervised the project. K.J.L. interpreted the results and finalized the paper. N-Y.S., G.H.K., and Y.H.H. performed the experiments and analyzed the data. S-K.K. provided brain tissue samples resected during surgery. J.Y.L provided neuropathological data. All authors participated in writing of the manuscript.

## Data availability

The data used for this study is available from the corresponding author upon reasonable request.

## Declarations

### Ethics Approval and Consent to Participate

Surgery was conducted following the acquisition of informed consent, in accordance with the Declaration of Helsinki. All procedures involving the use of human tissue received prior approval from the Institutional Review Board (Ethics Committee) of Seoul National University Medical Center (IRB No. H-0507-509-153).

### Consent to Publish declaration

Not applicable.

## References

1. Kabat J and Krol P (2012) Focal cortical dysplasia - review. Pol J Radiol 77(2): 35–43

2. Taylor DC, Falconer MA, Bruton CJ, and Corsellis JA (1971) Focal dysplasia of the cerebral cortex in epilepsy. J Neurol Neurosurg Psychiatry 34(4): 369–387

3. Kakita A, Kameyama S, Hayashi S, Masuda H, and Takahashi H (2005) Pathologic features of dysplasia and accompanying alterations observed in surgical specimens from patients with intractable epilepsy. J Child Neurol 20(4): 341–350

4. Roper SN and Yachnis AT (2002) Cortical dysgenesis and epilepsy. Neuroscientist 8(4): 356–371

5. Bronen RA, et al. (1997) Focal cortical dysplasia of Taylor, balloon cell subtype: MR differentiation from low-grade tumors. AJNR Am J Neuroradiol 18(6): 1141–1151

6. Mischel PS, Nguyen LP, and Vinters HV (1995) Cerebral cortical dysplasia associated with pediatric epilepsy. Review of neuropathologic features and proposal for a grading system. J Neuropathol Exp Neurol 54(2): 137–153

7. Palmini A, et al. (2004) Terminology and classification of the cortical dysplasias. Neurology 62(6 Suppl 3): S2-8

8. DeFelipe J, et al. (1993) Selective changes in the microorganization of the human epileptogenic neocortex revealed by parvalbumin immunoreactivity. Cereb Cortex 3(1): 39–48

9. Avoli M, et al. (2016) Specific imbalance of excitatory/inhibitory signaling establishes seizure onset pattern in temporal lobe epilepsy. J Neurophysiol 115(6): 3229–3237

10. Avoli M, et al. (2003) Epileptiform synchronization in the human dysplastic cortex. Epileptic Disord 5 Suppl 2: S45–50

11. 11. Duong T, De Rosa MJ, Poukens V, Vinters HV, and Fisher RS (1994) Neuronal cytoskeletal abnormalities in human cerebral cortical dysplasia. Acta Neuropathol 87(5): 493–503

12. Lurton D, et al. (2002) Immunohistochemical study of six cases of Taylor’s type focal cortical dysplasia: correlation with electroclinical data. Epilepsia 43 Suppl 5: 217–219

13. Mathern GW, et al. (2000) Neurons recorded from pediatric epilepsy surgery patients with cortical dysplasia. Epilepsia 41 Suppl 6: S162–167

14. Mizoguchi M, Iwaki T, Morioka T, Fukui M, and Tateishi J (1998) Abnormal cytoarchitecture of cortical dysplasia verified by immunohistochemistry. Clin Neuropathol 17(2): 100–109

15. Spreafico R, et al. (1998) Cortical dysplasia: an immunocytochemical study of three patients. Neurology 50(1): 27–36

16. Houser CR, Harris AB, and Vaughn JE (1986) Time course of the reduction of GABA terminals in a model of focal epilepsy: a glutamic acid decarboxylase immunocytochemical study. Brain Res 383(1-2): 129–145

17. Ribak CE (1985) Axon terminals of GABAergic chandelier cells are lost at epileptic foci. Brain Res 326(2): 251–260

18. Ribak CE, Hunt CA, Bakay RA, and Oertel WH (1986) A decrease in the number of GABAergic somata is associated with the preferential loss of GABAergic terminals at epileptic foci. Brain Res 363(1): 78–90

19. Sloviter RS (1991) Permanently altered hippocampal structure, excitability, and inhibition after experimental status epilepticus in the rat: the "dormant basket cell" hypothesis and its possible relevance to temporal lobe epilepsy. Hippocampus 1(1): 41–66

20. Calcagnotto ME, Paredes MF, Tihan T, Barbaro NM, and Baraban SC (2005) Dysfunction of synaptic inhibition in epilepsy associated with focal cortical dysplasia. J Neurosci 25(42): 9649–9657

21. Babb TL, et al. (2000) Brain plasticity and cellular mechanisms of epileptogenesis in human and experimental cortical dysplasia. Epilepsia 41 Suppl 6: S76–81

22. Ying Z, et al. (1999) Selective coexpression of NMDAR2A/B and NMDAR1 subunit proteins in dysplastic neurons of human epileptic cortex. Exp Neurol 159(2): 409–418

23. DeFelipe J, Huntley GW, del Rio MR, Sola RG, and Morrison JH (1994) Microzonal decreases in the immunostaining for non-NMDA ionotropic excitatory amino acid receptor subunits GluR 2/3 and GluR 5/6/7 in the human epileptogenic neocortex. Brain Res 657(1-2): 150-158

24. Marco P and DeFelipe J (1997) Altered synaptic circuitry in the human temporal neocortex removed from epileptic patients. Exp Brain Res 114(1): 1–10

25. Alonso-Nanclares L, et al. (2005) Microanatomy of the dysplastic neocortex from epileptic patients. Brain 128(Pt 1): 158–173

26. Rossini L, et al. (2021) Dendritic pathology, spine loss and synaptic reorganization in human cortex from epilepsy patients. Brain 144(1): 251–265

27. Redecker C, et al. (2005) Optical imaging of epileptiform activity in experimentally induced cortical malformations. Exp Neurol 192(2): 288–298

28. Du F, Eid T, Lothman EW, Kohler C, and Schwarcz R (1995) Preferential neuronal loss in layer III of the medial entorhinal cortex in rat models of temporal lobe epilepsy. J Neurosci 15(10): 6301–6313

29. Du F, et al. (1993) Preferential neuronal loss in layer III of the entorhinal cortex in patients with temporal lobe epilepsy. Epilepsy Res 16(3): 223–233

30. Nguyen HB, et al. (2016) Conductive resins improve charging and resolution of acquired images in electron microscopic volume imaging. Sci Rep 6: 23721

31. Gray EG (1959) Axo-somatic and axo-dendritic synapses of the cerebral cortex: an electron microscope study. J Anat 93: 420–433

32. Harvey CD and Svoboda K (2007) Locally dynamic synaptic learning rules in pyramidal neuron dendrites. Nature 450(7173): 1195–1200

33. Iascone DM, et al. (2020) Whole-Neuron Synaptic Mapping Reveals Spatially Precise Excitatory/Inhibitory Balance Limiting Dendritic and Somatic Spiking. Neuron 106(4): 566–578 e568

34. Sarnat HB and Flores-Sarnat L (2013) Radial microcolumnar cortical architecture: maturational arrest or cortical dysplasia? Pediatr Neurol 48(4): 259–270

35. Kubota Y, Karube F, Nomura M, and Kawaguchi Y (2016) The Diversity of Cortical Inhibitory Synapses. Front Neural Circuits 10: 27

36. Ikegaya Y, et al. (2001) Rapid and reversible changes in dendrite morphology and synaptic efficacy following NMDA receptor activation: implication for a cellular defense against excitotoxicity. J Cell Sci 114(Pt 22): 4083–4093

37. Swann JW, Al-Noori S, Jiang M, and Lee CL (2000) Spine loss and other dendritic abnormalities in epilepsy. Hippocampus 10(5): 617–625

38. Liu G (2004) Local structural balance and functional interaction of excitatory and inhibitory synapses in hippocampal dendrites. Nat Neurosci 7(4): 373–379

39. Mel BW and Schiller J (2004) On the fight between excitation and inhibition: location is everything. Sci STKE 2004(250): PE44

40. Devine MJ and Kittler JT (2018) Mitochondria at the neuronal presynapse in health and disease. Nat Rev Neurosci 19(2): 63–80

41. Vos M, Lauwers E, and Verstreken P (2010) Synaptic mitochondria in synaptic transmission and organization of vesicle pools in health and disease. Front Synaptic Neurosci 2: 139

42. Jedlicka P and Deller T (2017) Understanding the role of synaptopodin and the spine apparatus in Hebbian synaptic plasticity - New perspectives and the need for computational modeling. Neurobiol Learn Mem 138: 21–30

43. Maggio N and Vlachos A (2014) Synaptic plasticity at the interface of health and disease: New insights on the role of endoplasmic reticulum intracellular calcium stores. Neuroscience 281: 135–146

44. Kato D and Wake H (2019) Activity-Dependent Myelination. Adv Exp Med Biol 1190: 43–51

45. Noori R, et al. (2020) Activity-dependent myelination: A glial mechanism of oscillatory self-organization in large-scale brain networks. Proc Natl Acad Sci U S A 117(24): 13227–13237

46. Knowles JK, et al. (2022) Maladaptive myelination promotes generalized epilepsy progression. Nat Neurosci 25(5): 596–606

47. Citraro R, Leo A, Constanti A, Russo E, and De Sarro G (2016) mTOR pathway inhibition as a new therapeutic strategy in epilepsy and epileptogenesis. Pharmacol Res 107: 333–343

48. Jansen LA, et al. (2015) PI3K/AKT pathway mutations cause a spectrum of brain malformations from megalencephaly to focal cortical dysplasia. Brain 138(Pt 6): 1613–1628

49. Weckhuysen S, et al. (2016) Involvement of GATOR complex genes in familial focal epilepsies and focal cortical dysplasia. Epilepsia 57(6): 994–1003

50. Multani P, Myers RH, Blume HW, Schomer DL, and Sotrel A (1994) Neocortical dendritic pathology in human partial epilepsy: a quantitative Golgi study. Epilepsia 35(4): 728–736

51. Obi-Nagata K, et al. (2023) Distorted neurocomputation by a small number of extra-large spines in psychiatric disorders. Sci Adv 9(23): eade5973

52. Tatti R, Haley MS, Swanson OK, Tselha T, and Maffei A (2017) Neurophysiology and Regulation of the Balance Between Excitation and Inhibition in Neocortical Circuits. Biol Psychiatry 81(10): 821–831

