## Supplementary material for "Synaptic Ultrastructural Alterations in Human Focal Cortical Dysplasia: Insights from Volume Electron Microscopy": Supplemntal Figures


### Supplementary Data


Supplementary Fig. 1. Histopathological Assessment of the Epileptogenic Cortex.

Supplementary Fig. 2. Reduction of Inhibitory Synaptic Density in Pyramidal Neuron Somata.

Supplementary Fig. 3. 3D Reconstruction of Distal Dendritic Segments of Pyramidal Neurons Using SB-SEM.

Supplementary Fig. 4. Abnormal Enlargement of Dendritic Spines in the Epileptogenic Region.

Supplementary Fig. 5. Enlargement Excitatory Synapses with More Vesicles in the Epileptogenic Region.

Supplementary Fig. 6. Schematic Synaptic Alterations in the FCD Cortex.

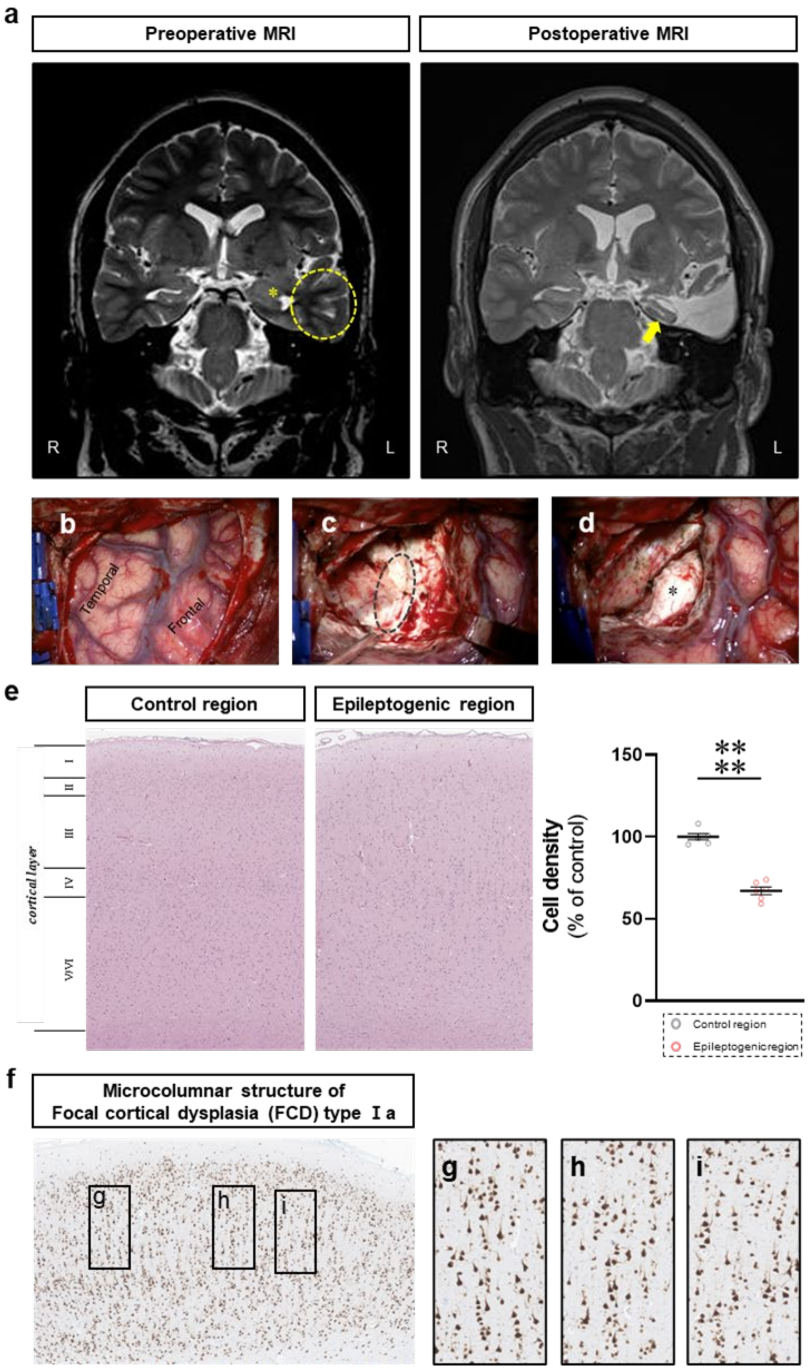


**Supplementary Fig. 1. Histopathological Assessment of the Epileptogenic Cortex.** Representative coronal T2-weighted MRI images **(a)**. Preoperative MRI shows blurring of the gray-white matter discrimination (*) and decreased subcortical white matter signal (dotted circle) in the anteromedial aspect of the left inferior temporal lobe. Postoperative MRI reveals anterior temporal lobectomy and resection of the parahippocampal gyrus, with the hippocampus (arrow) preserved. R: right, L: left. Intraoperative photographs **(b-d)**. Left temporal and frontal lobes are exposed after craniotomy and durotomy (b). The margin between the grey and white matter (dotted circle) in the inferior temporal lobe is visible following temporal lobectomy (c). Hippocampus (*) is shown after lesion resection (d). H&E staining **(e)**. Note the reduced cell density in cortical layer III of the epileptogenic area, without dysmorphic neurons or balloon cells. ****$P$ < 0.0001. Data are represented as mean ± SEM. NeuN staining **(f)**. Note a characteristic microcolumnar arrangement of neurons, confirming focal cortical dysplasia type Ia **(g-i)**.

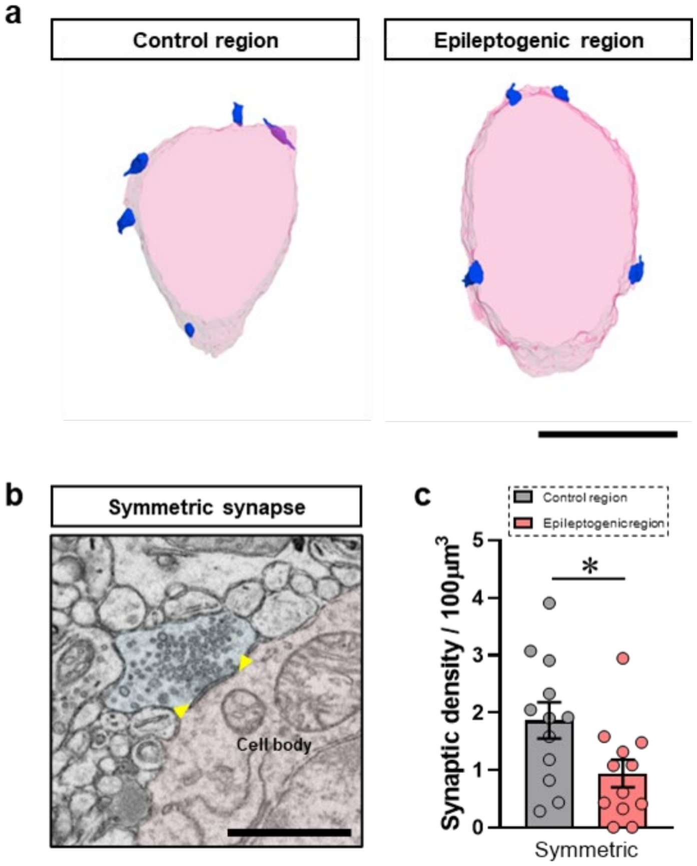


**Supplementary Fig. 2. Reduction of Inhibitory Synaptic Density in Pyramidal Neuron Somata.** Representative 3D reconstruction images of pyramidal neuron cell bodies displaying somatic inhibitory synapses **(a)**. Representative ATUM-SEM image showing an inhibitory (symmetric) synapse on the cell body, with yellow arrows highlighting the postsynaptic density (PSD) **(b)**. Quantification of somatic inhibitory synapse density **(c)**. *$P < 0.05$; $n = 12$ per group; unpaired Student's t-test. Data are presented as mean ± SEM.

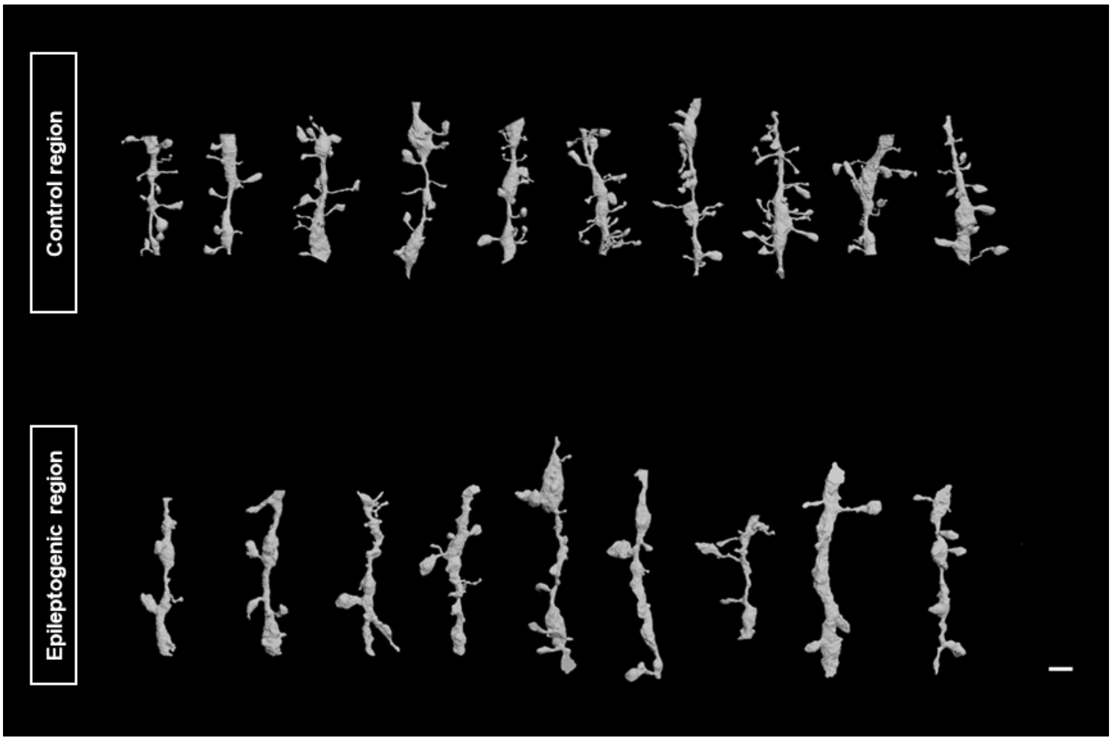


**Supplementary Fig. 3. 3D Reconstruction of Distal Dendritic Segments of Pyramidal Neurons Using SB-SEM.** Representative 3D reconstruction images of distal dendritic segments from cortical layer III pyramidal neurons, showing dendritic spines in the control (temporal cortex) and epileptogenic (uncus) regions. Note the presence of a subset of extra-large spines in the epileptic area.

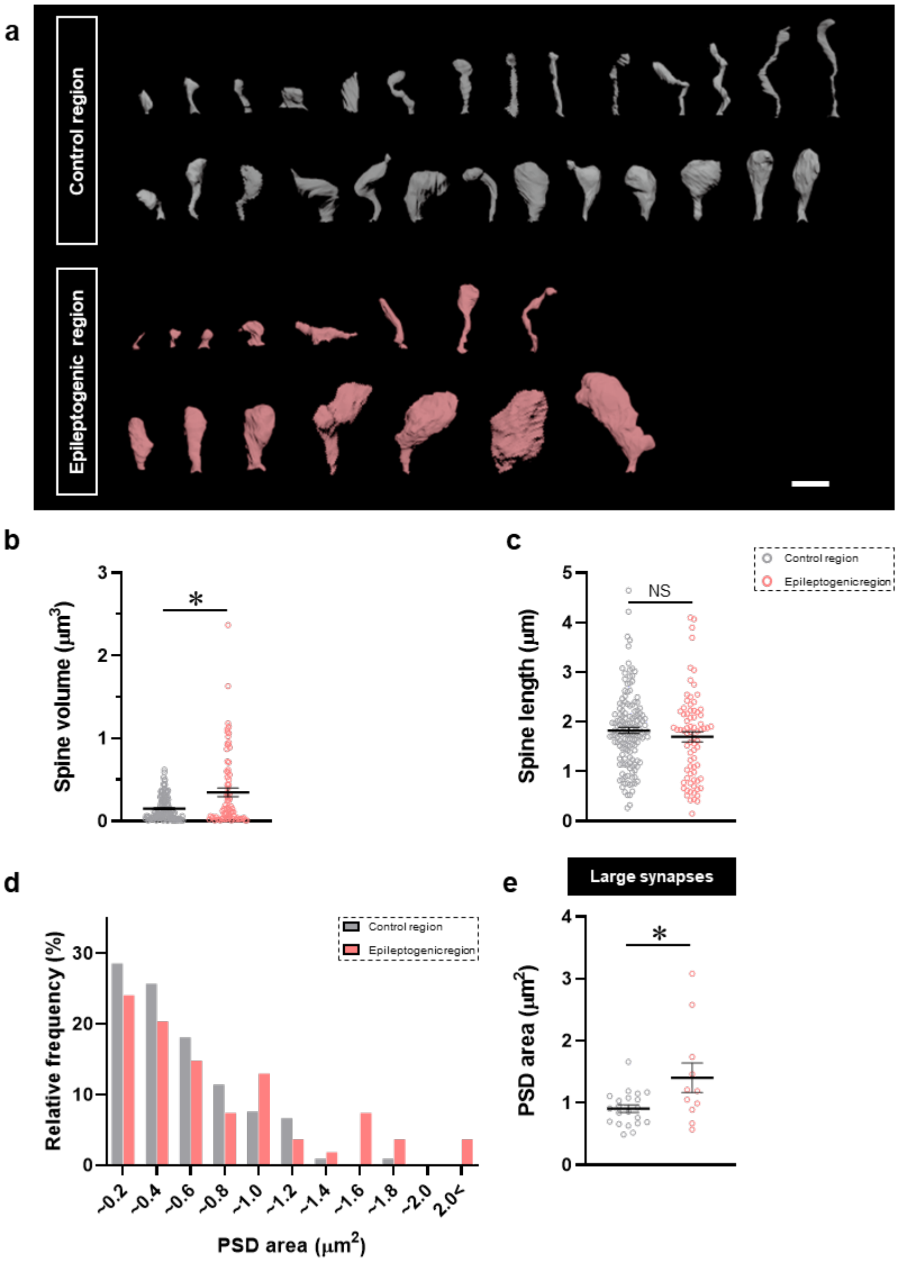


**Supplementary Fig. 4. Abnormal Enlargement of Dendritic Spines in the Epileptogenic Region.** Representative image of dendritic spine shapes in the control (gray) and epileptogenic (pink) regions **(a)**. Scale bar, 1 μm. Spines from two dendritic segments are arranged by size. Mean spine volumes **(b)** and spine lengths **(c)** measured from the dendrite surface to the spine tip (control, n = 150; epilepsy, n = 76). NS, not significant. *P<0.05; Mann-Whitney U test. Cumulative distribution of PSD areas (control, n = 105; epilepsy, n = 54) **(d)**. Average PSD areas in the top 20% of the population in D (control, n = 21; epilepsy, n = 11) **(e)**. *P < 0.05; unpaired Student's t-test. Data are presented as mean ± SEM.

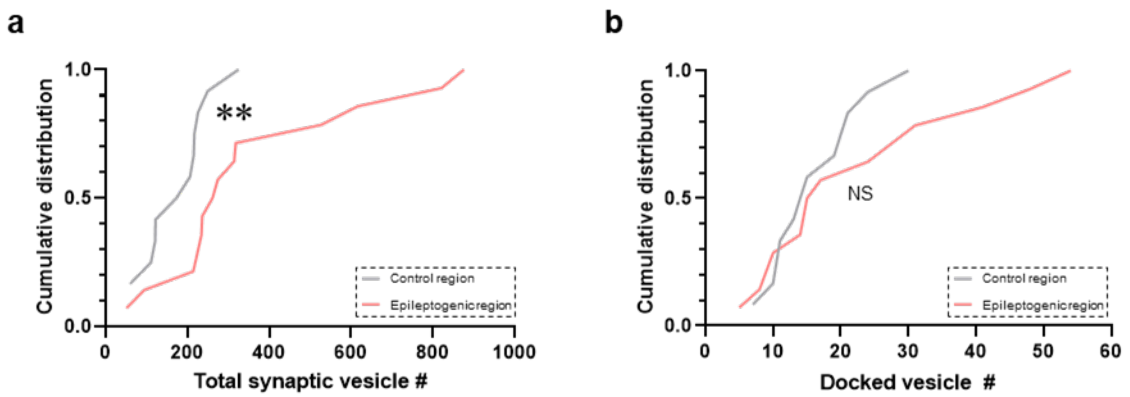


**Supplementary Fig. 5. Enlargement Excitatory Synapses with More Vesicles in the Epileptogenic Region.** Cumulative distributions of total **(a)** and docked **(b)** synaptic vesicles in individual presynaptic boutons (n = 12-14 boutons/region). NS, not significant. **P<0.01; Kolmogorov-Smirnov test.

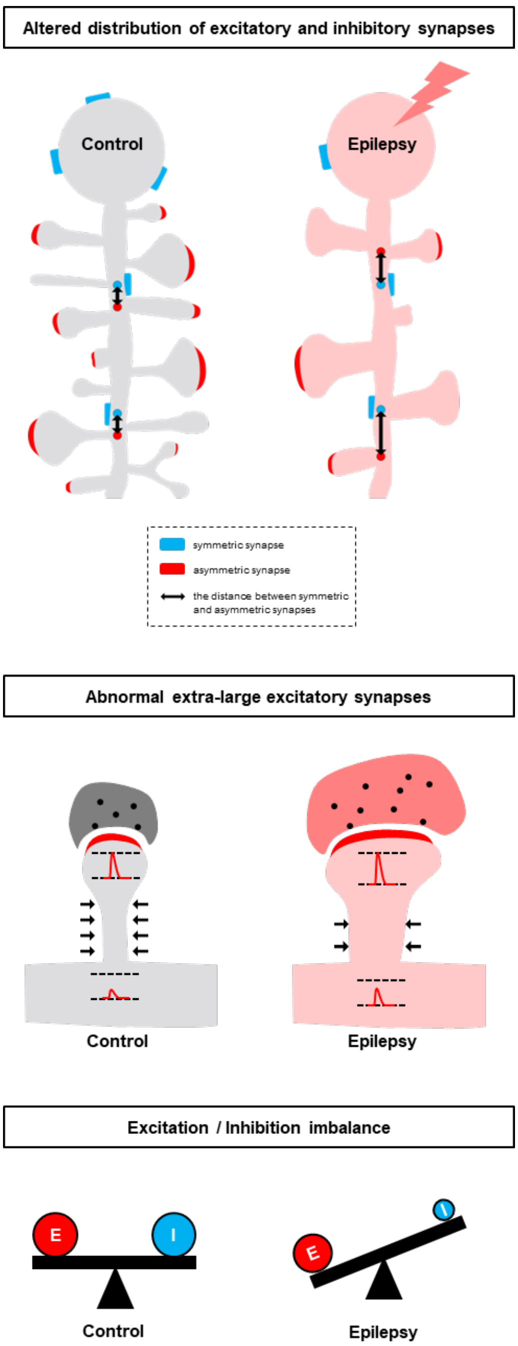


**Supplementary Fig. 6. Schematic Model of Synaptic Alterations Underlying Hyperexcitability in the FCD Cortex.** Key findings of our volume electron microscopy study include (1) a reduced number of inhibitory synapses on pyramidal neuron somata, (2) the presence of extra-large excitatory synapses with increased presynaptic vesicles, (3) an elongated distance between inhibitory and excitatory synapses, and (4) maladaptive myelination (not shown here; see **Fig. 5**). Collectively, these structural changes suggest that multiple mechanisms can contribute to neuronal hyperexcitability and seizure propagation in FCD.
